# Migratory pattern of zoonotic *Toxocara cati* and *T. canis* in experimentally infected pigs

**DOI:** 10.1101/2023.04.27.538522

**Authors:** Casper Sahl Poulsen, Ayako Yoshida, Tinna Thordardottir Wellbrant, Pall Skuli Leifsson, Per Skallerup, Stig Milan Thamsborg, Peter Nejsum

**Affiliations:** Herlev and Gentofte Hospital, 2820 Copenhagen, Denmark; University of Miyazaki, Miyazaki, Japan; University of Copenhagen, Copenhagen, Denmark; Statens Serum Institut, Copenhagen, Denmark; Aarhus University and Aarhus University Hospital, Aarhus, Denmark

**Keywords:** *Toxocara canis*, *Toxocara cati*, zoonosis, migration pattern, pathology, histopathology

## Abstract

Over a billion people are infected with *Toxocara canis* or *T. cati*, the roundworms of dogs and cats. Historically, *T. canis* has been considered the main responsible of human toxocarosis but as serodiagnosis cannot discriminate the two species, this remains unresolved. We assessed the migratory pattern of *T. cati* and *T. canis* in a pig model and found them to be equally infective. Overall, they had a similar migration pattern reaching multiple organs and tissues, including mesenteric lymph nodes, liver, lungs and diaphragm. We recovered larvae of both species in the brain, suggesting that *T. cati* also can cause neurological toxocarosis in humans. Both species induced systemic eosinophilia and histopathological changes in lungs, livers and mesenteric lymph nodes. This study emphasizes the importance of *T. cati* as a zoonotic agent and the need to develop diagnostic methods that can differentiate between sources of infection in humans.

## Introduction

*Toxocara canis* and *T. cati* are common roundworms parasitizing dogs and cats, respectively. The global prevalence of *T. cati* is estimated to be 17.0% in approximately 118-150 million cats worldwide, while it is 11.1% for *T. canis* in the ≥100 million dogs (1,2). Ingestion of infective eggs can cause disease in humans due to larvae migration, and it has recently been estimated that 1.2 billion humans are exposed to or are infected with *Toxocara* spp. Therefore, toxocarosis is now listed as one of the five parasitosis prioritised for public health action in the US by the Centers for Disease Control (CDC) (3) and is also highly prioritized in Europe (4,5). Symptoms depend on infection dose, the route of larval migration, frequency of reinfection and the host response (6). Systemic migration of larvae is termed Visceral Larva Migrans (VLM) (7). Larval invasion of the eye was described two years earlier (8) and later termed Ocular Larva Migrans (OLM). Less severe clinical manifestation has been classified as covert toxocarosis in children (9) and common toxocarosis in adults (10). These are probably the same syndromes with variation in relation to age (11). A fifth syndrome where larvae migrate in the CNS is termed Neurological Toxocarosis (NT) (6), which may cause epilepsy as an association between *Toxocara* spp. serum antibodies and seizures have recently been observed (12). The impact of NT was also recently investigated (13) where infection was associated with neurodegeneration and major alteration in the transcriptional profile in the brains of mice.

The relative importance of *T. canis* and *T. cati* associated with human disease is an ongoing discussion but historically far most attention has been given to *T. canis* (14–18), despite that no widely available serologic diagnostic method can distinguish between the two parasitic infections in humans (15,17–19). The zoonotic potential and consequences for human health of *T. canis* and *T. cati* have been explored by investigating the migratory behavior of the parasite and the associated pathological changes in the affected organs in experimental animal models (14,16). Experimental infections of mongolian gerbils indicate that *T. canis* larvae have higher affinity for the eyes than *T. cati* (20), and studies in mice suggest that *T. canis* larvae accumulate in the brain whereas *T. cati* accumulate in the muscle tissue (21). The pig has been suggested to be a good model for human toxocarosis due to similar size, weight, immune response, liver physiology and metabolic function (22–24). Experimental studies in the pig assessing the migratory pattern and associated pathology of *T. canis* (25–33), and *T. cati* (34) have reported larval recoveries from a variety of organs and muscles, including lymph nodes, liver, lungs, eyes, kidneys, diaphragm, tongue, and masseter. While the large majority of the experimental studies in pigs have focused on either *T. canis* or *T. cati*, only a single comparative study has been performed. The author found that *T. cati* migrates to the lymph nodes, livers and lungs in pigs; however, the number of larvae was not quantified and only macro- and microscopic changes in the liver were assessed (26).

The objectives of the present study were to compare the migratory behaviour of *T. canis* and *T. cati* larvae and associated pathological changes in pigs. The migratory behaviour was evaluated by recovery of larvae from different organs after necropsy. The pathological assessment comprised histopathology to investigate the inflammatory response and fibrosis in the liver, lungs and mesenteric lymph nodes, enumeration of white spots on livers and kidneys, and blood eosinophilia.

## Materials and methods

### Experimental design and animals

Two different experimental infection studies were conducted and are termed experiment 1 (Exp. 1) and experiment 2 (Exp. 2), respectively.

Exp. 1: Seventeen helminth-naïve Danish Landrace/Yorkshire/Duroc crossbred pigs eight weeks of age (body weight range: 17-28; mean: 22.4 kg) were obtained from a commercial specific pathogen free (SPF) breeder. The pigs were ear tagged and allocated into three groups after stratification according to sex and weight. In groups 1 (n=6) and 2 (n=5) pigs were inoculated by stomach tube with a single dose of 50,000 embryonated *T. canis* and *T. cati* eggs, respectively. Group 3 (n=6) served as uninfected controls inoculated with tap water. Pigs were necropsied at 14 days post infection (dpi).

Exp. 2: Performed as Exp. 1, but the infective dose of both *T. canis* and *T. cati* was 10,000 embryonated eggs and pigs were necropsied at 31 dpi (due to logistic reasons, these pigs were necropsied over three days (30, 31 and 32 dpi)). The infected groups and the control group included seven and four pigs, respectively. All males in both experiments were castrated.

EDTA-stabilised blood samples were obtained at days 0, 7 and 14 dpi in Exp. 1 to evaluate the numbers of eosinophil granulocytes. Samples were analysed the same day at the Central Laboratory at the Faculty of Medical and Health Sciences, the University of Copenhagen, Denmark.

All pigs were fecal negative for *Ascaris suum* eggs 0 dpi (35) and seronegative to *T. canis* and *A. suum* antibodies (36) before experimental infection.

### Housing, infective material and study approval

The pigs were housed in three separate rooms that had been thoroughly washed, and flame-cleaned prior to use. Separate boots, protective overalls and tools were used for each group. The pigs were fed a standard diet consisting of ground barley with a protein/mineral supplement and ad libitum access to water and allowed to acclimatize for one week before inoculation.

Embryonated *T. canis* and *T. cati* eggs were kindly provided by colleagues at University of Veterinary Medicine Hannover, Germany, and stored in 0.05 M H_2_SO_4_ at five degrees until use. The viability of the eggs was tested in a hatching assay and found to be similar for *T. canis* and *T. cati* (80-90 %).

Larvae isolated from *T. canis* (n=5) and *T. cati* (n=5) infected pigs (see below) had their partial ITS1 and complete 5.8s rRNA gene and ITS2 Sanger sequenced (37) and found to be 100% identical to the sequence of *T. canis* (OM876369.1) and 99.85-100 % identical to *T. cati* (KY003086.1) (GenBank accession numbers: LC762618 - LC762621).

The studies were approved by the Animal Experiments Inspectorate, Ministry of Justice, Denmark (Ref. 2010/561-1914).

### Necropsy and processing of organs

Exp. 1: The pigs were necropsied on day 14 dpi using a captive bolt pistol to stun the pigs followed by exsanguination. The liver (without gallbladder), lungs, mesenteric lymph nodes (MLN), brain, eyes, and body muscles (pooling 100 g of front limb, hind limb, and loin) were sampled from the infected pigs while only the liver, lungs and MLN were included for the control pigs.

White spots on liver and kidney surfaces were counted and for the liver, the white spots were identified as either granulation-tissue type or lymphonodular (26). Before further processing subsamples were taken for histological examination (see below).

The digestion of organs and counting of larvae was conducted according to (30). Briefly, organ weights were noted and blended in a food processor until a tissue fragments a size of 2-3 mm^3^. If organ weight was >100g subsampling was used. All samples were digested with HCl/pepsin at 45 °C for 60 minutes under continuous stirring. Then three sedimentation steps (30 minutes each) were performed and samples stored in 70 % ethanol at 5 °C until enumeration.

Exp. 2: Slaughtering, processing of organs/tissues and counting were carried out the same way as in Exp. 1, but the heart, diaphragm and tongue were also included in the analysis.

All larvae counts were converted into the total number of larvae in the whole organ, except for larvae recovered from muscles.

### Histology and haematology

Samples were taken from the liver, left lung and MLN on 14 and 31 dpi. Tissues were fixed in 4% formaldehyde in phosphate-buffered saline. After dehydration, samples were embedded in paraffin, sectioned at 2-4 μg and stained with hematoxylin and eosin (HE). Eosinophilia was categorized after the number of eosinophils: none-mild (<50 per high-power field) or moderate-massive (≥50). To facilitate the evaluation of fibrosis, selected slides were stained for connective tissue by the Masson Trichrome (MT) technique (38), and fibrosis was graded as nil-mild, moderate or massive, as defined in Appendix Figure 1-2. Selected samples were immunohistochemically stained for ionized calcium-binding adaptor molecule 1 (IBA1) to identify macrophages and facilitate evaluation of the inflammatory response. An Avidin/Biotin Complex (ABC) method was used, where non-specific binding cites were blocked with 4% normal rabbit serum (X0902; Dako, DK), the primary antibody was a polyclonal goat anti-IBA1 (ab5076; Abcam, UK), and the secondary antibody was a biotinylated rabbit anti-goat (E0466; Dako, DK) (39).

**Figure 1.**
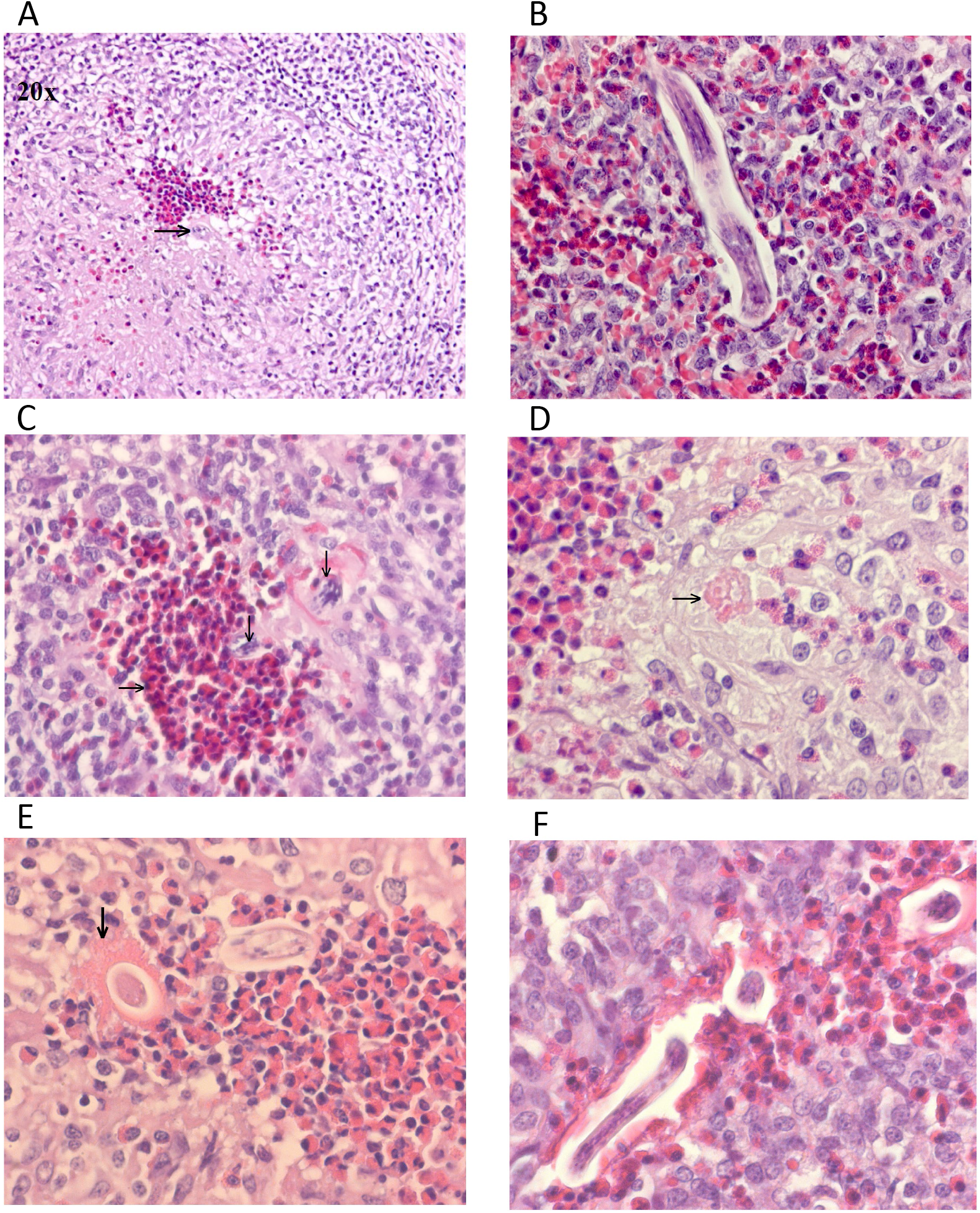
A: Lung granuloma, from a *Toxocara canis* infected pig (31 dpi, 10,000 eggs). A larva (→) and eosinophils in center surrounded by macrophages, lymphocytes, and fibroblasts (obj. X 20) B: Larva in the center of a lung granuloma from a *T. canis* infected pig (14 dpi, 50,000 eggs) (obj. X 40) C: Eosinophils (→) and larvae (↓) in the center of a lung granuloma from a *T. cati* infected pig (31 dpi, 10,000 eggs) (obj. X 40) D: Center of a mesentery lymph node (MLN) granuloma, from a *T. canis* infected pig (31 dpi, 10,000 eggs) with an eosinophilic granular mass (→) that possibly represents a larva residue (obj. X 60) E: Center of a MLN granuloma in a *T. cati* infected pig (31 dpi., 10,000 eggs). A larva surrounded by a flaming eosinophilic mass (↓), eosinophils, and macrophages (obj. X 60) F: Center of a MLN granuloma in a *T. cati* infected pig (14 dpi., 50,000 eggs) with larvae surrounded by eosinophils and macrophages (obj. X 60)

In Exp. 1 standard haematological analyses were performed using an ADVIA2120 haematology analyser (Siemens), including white blood cells (WBC) and eosinophils (EOS).

### Statistics

Statistical analysis was performed in R (v4.2.0). Comparison of larvae counts, white spots and eosinophil levels between infected groups were performed non-parametrically since hypotheses of normality were rejected (Shapiro-Wilk). Kruskal-Wallis test was used to evaluate the effect of the group and, if significant, followed by pairwise comparison using a Mann-Whitney test. Visualization of eosinophil levels was performed with ggplot2 (v3.3.5). Fisher exact test was used to compare the histological eosinophilia grading and fibrosis score of *T. canis* and *T. cati* infected pigs. *P*-value of less than 0.05 was considered statistically significant.

## Results

### Migratory pattern

Two weeks after infection the large majority of larvae were found in the MLN and lungs for both species. The median total number of recovered larvae from each pig was 145 and 70 for *T. canis* and *T. cati*-infected pigs, respectively in Exp. 1 (*P* = 0.27) (Table 1). No statistical differences in recoveries of *T. canis* and *T. cati* were found for any of the individual organs/tissues. It is noted that three of the *T. canis* infected pigs had larvae in the livers while none were found in the *T. cati* infected pigs.

**Table 1.** The number of larvae recovered at 14 dpi. and at 31 dpi. from the mesenteric lymph nodes (MLN), liver, lungs, brain and in total from pigs infected with 50,000 and 10,000 *Toxocara* spp. eggs, respectively. Pigs are listed in the same order.

| Organ | 14 dpi. (50,000 eggs) |  |  |  |  | 31 dpi. (10,000 eggs) |  |  |  |  |
| --- | --- | --- | --- | --- | --- | --- | --- | --- | --- | --- |
|  | *T. canis* (n=6) |  | *T. cati* (n=5) |  | *P*-value* | *T. canis* (n=7) |  | *T. cati* (n=7) |  | *P*-value |
|  | Larvae counts | Median | Larvae counts | Median |  | Larvae counts | Median | Larvae counts | Median |  |
| MLN | 88,53,165,19,71,203 | 80 | 21,48,161,384,57 | 57 | 0.86 | 1,3,2,1,1,0,1 | 1 | 9,2,0,2,5,3,4 | 3 | 0.06 |
| Liver | 0,10,0,65,83,0 | 5 | 0,0,0,0,0 | 0 | 0.08 | 0,0,0,0,0,11,0 | 0 | 0,0,0,0,0,0,0 | 0 | n.a. |
| Lungs | 5,24,0,38,421,86 | 31 | 30,21,86,149,0 | 30 | 1 | 12,16,12,0,32,0,4 | 12 | 0,8,0,3,0,3,0 | 0 | 0.07 |
| Brain | 7,1,0,2,0,3 | 2 | 1,1,1,0,0 | 1 | 0.29 | 0,0,0,0,0,0,0 | 0 | 0,0,0,0,0,0,0 | 0 | n.a. |
| Total | 100,88,165,124,575,29<br>2 | 145 | 52,70,248,533,57 | 70 | 0.27 | 13,19,14,1,33,11,<br>5 | 13 | 9,10,0,5,5,6,4 | 5 | 0.06 |

\* Mann-Whitney test for *T. canis* vs. *T. cati*

Overall, we found lower recoveries on 31 dpi than on 14 dpi. In Exp. 2, none of the pigs had larvae in the brain and only one pig (*T. canis* group) had larvae in the liver. The same number of *T. canis* and *T. cati* were recovered overall and when comparing the individual organs/tissues; however, there was a tendency for a higher total recovery of *T. canis* than *T. cati* (median: 13 vs. 5; *P* = 0.06). Furthermore, there was a trend for more *T. cati* larvae in the MLN compared with *T. canis* larvae (median: 3 vs. 1; *P* = 0.06) while the opposite trend was seen in the lungs, where more larvae were recovered from the *T. canis-*infected pigs (*P* = 0.07) (Table 1). One larva from the diaphragm of a *T. canis* infected pig, and one larva from both diaphragm and body muscle samples in a *T. cati* infected pig was recovered at 31 dpi.

No larvae were recovered from the eyes in the two experiments. No larvae were found in the control pigs.

### White spots

The total number of liver white spots was similar in both infected groups at 14 dpi (*P* = 0.86) (Table 2). However, more lymphonodular liver white spots were observed on livers from *T. canis* infected pigs (*P* = 0.006) whereas more granulation-tissue type were observed for *T. cati* infected pigs (*P* = 0.045). A tendency for higher number of white spots on kidneys of *T. cati* infected pigs was observed at 14 dpi (*P* = 0.10).

**Table 2.** Number of white spots (WS) observed at 14 dpi. and at 31 dpi. on livers in total and given as diffuse WS (granulation-tissue) and lymphonodular WS and in total on kidneys from pigs infected with 50,000 and 10,000 *Toxocara* spp. eggs, respectively. Pigs in groups are listed in the same order.

| Organ | 14 dpi. |  |  |  |  | 31 dpi. |  |  |  |  |
| --- | --- | --- | --- | --- | --- | --- | --- | --- | --- | --- |
|  | *T. canis* (n=6) |  | *T. cati* (n=5) |  | *P*-value* | *T. canis* (n=7) |  | *T. cati* (n=7) |  | *P*-value |
|  | WS counts | Median | WS counts | Median |  | WS counts | Median | WS counts | Median |  |
| Liver: Granulation-tissue type | 373,417,261, 279,331,273 | 305 | 402,488,535, 580,292 | 488 | 0.045 | 446,293,71,291, 334,226,95 | 291 | 9,34,6,1,31, 31 | 20 | 0.003 |
| Liver: Lymphonodular type | 105,232,50, 215,231,63 | 160 | 5,5,6, 5,10 | 5 | 0.006 | 58,27,32,44, 32,3,8 | 32 | 6,1,0,0,1,3 | 1 | 0.005 |
| Liver: Total | 478,649,311, 494,562,336 | 486 | 407,493,541, 585,302 | 493 | 0.86 | 504,320,103,335, 366,229,103 | 320 | 15,35,6,1,32, 34 | 24 | 0.003 |
| Kidney: Total | 61,88,45, 57,64,38 | 59 | 1,190,224, 427,162 | 190 | 0.10 | 6,17,8,16, 13,2,3 | 8 | 0,0,4,5,1,1 | 1 | 0.015 |

\* Mann-Whitney test for *T. canis* vs. *T. cati*

At 31 dpi significantly higher numbers of white spots were observed on the livers (both types) and kidneys of *T. canis* infected pigs compared with *T. cati* infected pigs (Table 2).

The higher number of lymphonodular white spots on *T. canis* infected livers gave them a much more rugged appearance as compared to the *T. cati* livers. At 14 dpi the high number of white spots made the livers of both *T. canis* and *T. cati* infected pigs look greyish whereas at 31 dpi the white spots had reduced in size, and the livers had lost the greyish appearance looking normally reddish brown, similar to the control livers (Appendix Figure 3).

There were no white spots on the livers and kidneys of any control pigs at the two time-points.

### Histology

Eosinophilia was present at various extents and locations in the liver (perilobular, portal and interlobular), lungs (e.g. alveolar/interlobular septum, peribronchial, and pleura) and MLN (e.g. peripheral medulla, trabeculae and paracortex) of *T. canis* and *T. cati* infected pigs (Table 3, Figure 1, Appendix Table 1). Two and three of the non-infected control pigs presented with moderate eosinophilia in the MLN at 14 and 31 dpi, respectively. There were no differences between eosinophilia in the three organs of *T. canis* and *T. cati* infected animals at any of the two time-points, although a tendency for more eosinophilia in the liver and lungs of *T. canis* pigs was observed.

**Table 3.** Number of pigs presenting with none-mild and moderate-massive eosinophilia in liver, lungs and mesenteric lymph nodes (MLN) 14 dpi and 31 dpi. The pigs were infected with 50,000 and 10,000 *Toxocara* spp. eggs, respectively.

| 14 dpi. |  |  |  |  |  | 31 dpi. |  |  |  |  |
| --- | --- | --- | --- | --- | --- | --- | --- | --- | --- | --- |
| *T. canis* (n=6) |  |  | *T. cati* (n=5) |  |  | *T. canis* (n=7) |  | *T. cati* (n=7) |  |  |
| Organ | None-Mild | Moderate-Massive | None-Mild | Moderate-Massive | *P*-value * | None-Mild | Moderate-Massive | None-Mild | Moderate-Massive | *P*-value |
| Liver | 0 | 6 | 1 | 4 | 0.45 | 5 | 2 | 7 | 0 | 0.46 |
| Lungs | 0 | 6 | 0 | 5 | 1.00 | 5 | 2 | 6 | 1 | 1.00 |
| MLN | 0 | 6 | 0 | 5 | 1.00 | 0 | 7 | 0 | 7 | 1.00 |

\* Fisher exact test for *T. canis* vs. *T. cati*

A granulomatous reaction was observed in the liver of a *T. canis* infected pig 14 dpi and in another on 31 dpi. No granulomas were found in the liver of *T. cati* infected pigs or in the controls. Granulomas that sometimes included larvae and/or necrosis, were found in the lungs of four *T. canis* and three *T. cati* infected, and in the MLN of two *T. canis* and four *T. cati* infected at 14 dpi. Similar granulomas were found in the lungs of two *T. canis* and one *T. cati* infected and in the MLN of two *T. canis* and four *T. cati* infected pigs at 31 dpi (Figure 1).

Focal inflammatory reaction, consisting mainly of lymphocytes, was seen in the lung of one *T. canis* infected pig (at 31 dpi) and two *T. cati* infected pigs (at 14 and 31 dpi).

Fibrosis was present at variable extents and locations in the liver (portal and/or interlobular) and lungs (mainly interlobular) of *T. canis* and *T. cati* infected pigs (Table 4). Only one *T. canis* infected pig presented with fibrosis in the MLN at 14 dpi. Fibrosis was not observed in any of the control pigs at 14 and 31 dpi.

**Table 4.** Number of pigs presenting with liver- and lung fibrosis in *Toxocara* spp. infected pigs 14 dpi and 31 dpi. The pigs were infected with 50,000 and 10,000 *Toxocara* spp. eggs, respectively.

| 14 dpi. |  |  |  |  |  |  |  | 31 dpi. |  |  |  |  |  |  |  |
| --- | --- | --- | --- | --- | --- | --- | --- | --- | --- | --- | --- | --- | --- | --- | --- |
| *T. canis* (n=6) |  |  |  | *T. cati* (n=5) |  |  |  | *T. canis* (n=7) |  |  |  | *T. cati* (n=7) |  |  |  |
| Organ | None | Moderate | Massive | None | Moderate | Massive | *P*-value* | None | Moderate | Massive |  | None | Moderate | Massive | *P*-value |
| iver | 1 | 1 | 4 | 2 | 3 | 0 | 0.11 | 3 | 1 | 2 |  | 7 | 0 | 0 | 0.19 |
| Lungs | 4 | 2 | 0 | 2 | 2 | 1 | 0.74 | 4 | 3 | 0 |  | 5 | 2 | 0 | 1.00 |

\* Fisher exact test for *T. canis* vs. *T. cati*

All infected pigs in Exp. 1 had blood eosinophilia seven- and 14-days dpi, and no significant difference in levels between *T. canis* and *T. cati* infected pigs was observed (Figure 2). The eosinophilia was also reflected in higher counts of WBC in infected groups (data not shown).

**Figure 2.**
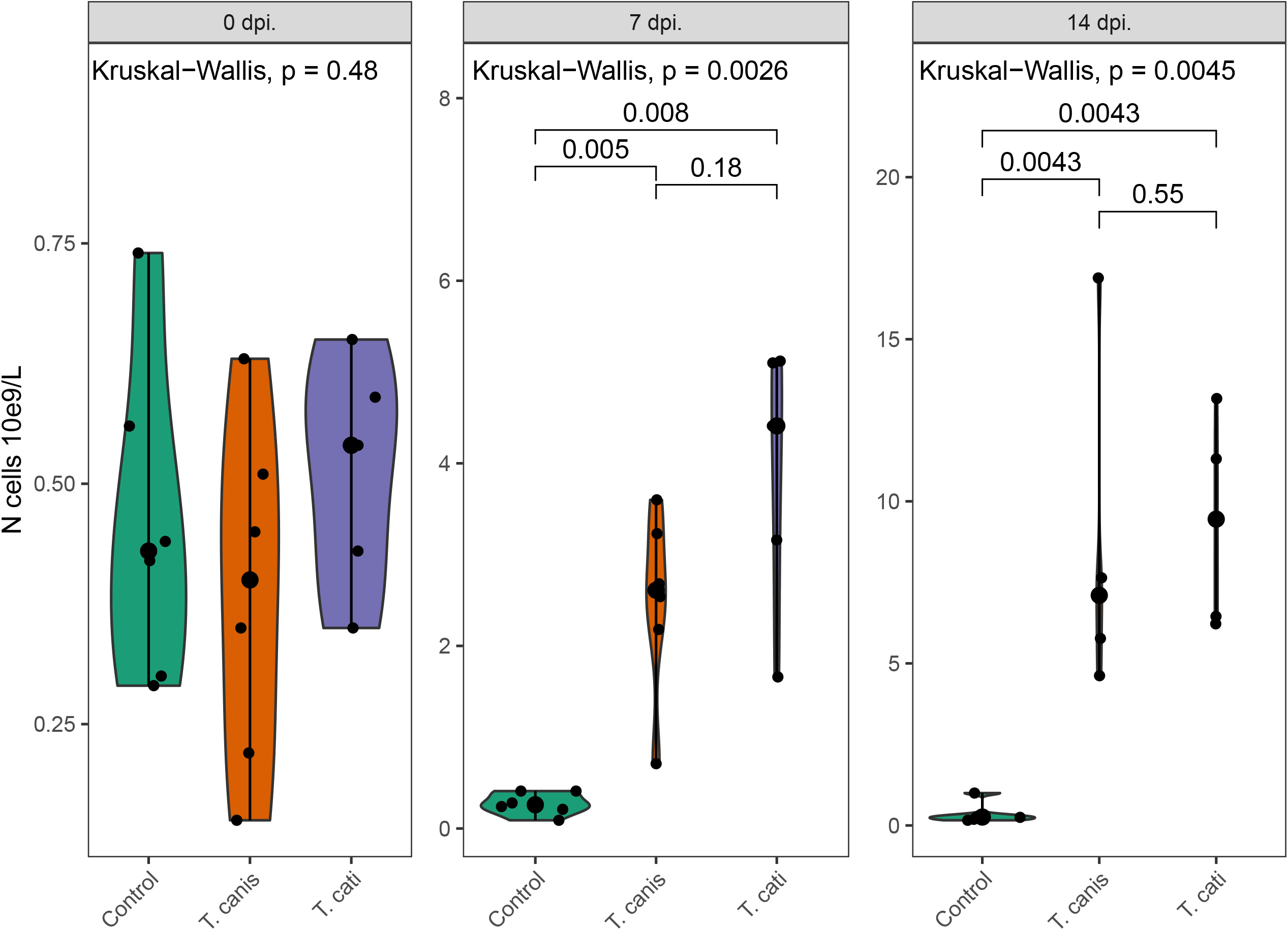
Eosinophil levels in the blood of pigs infected with 50,000 infective *Toxocara canis* or *T. cati* eggs or uninfected controls at days 0, 7 and 14 dpi. Each dot represents one pig and the enlarged dot the median. The group effect was tested with a Kruskal-Wallis test at each timepoint and if significant pairwise comparison was performed with Mann-Whitney U tests.

## Discussion

The zoonotic potential of *T. canis* has been acknowledged for decades whereas *T. cati* is most often ignored probably due to diagnostic issues in humans and a lack of experimental evidence in larger animals (15). We therefore compared the migratory capacity and associated pathology of *T. cati* and *T. canis* using the pig as a model for human infection. Two weeks after infection, we found similar total numbers of *T. cati* and *T. canis* larvae, suggesting that *T. cati* is as infective to pigs as the well-studied *T. canis*. This observation was consistent for all organs examined. Despite initial similarities between the two species, we found suggestive evidence for different migratory patterns later in the infection (day 31 dpi), with a tendency for more *T. cati* larvae in the MLN whereas more *T. canis* was found in the lungs (Table 1). In addition, *T. canis* persisted in the liver for longer time and caused more granulation-tissue type white spots than *T. cati*.

The specific migration route of *Toxocara* spp. in pigs is unknown, and larvae seem to find their way to most organs of the host. Previous studies have shown that *T. canis* larvae are found in the MLN and livers 14 dpi, with numbers peaking in the lungs one week later while similar larval numbers were found in lymph nodes and lungs at day 7 and 14 dpi in *T. cati* infected pigs (27,30,31,34). In contrast to *T. canis*, we recovered no *T. cati* from the livers at 14 dpi despite high numbers of white spots, suggesting that the larvae have left this organ at the time of necropsy. In accordance with a previous study where none to very few *T. cati* larvae in the livers of pigs infected with 100,000 eggs (34). The high number of lymphonodular white spots and tendency for more fibrosis in the livers of *T. canis* infected pig (see below) may retain the larvae in the livers and may therefore explain the difference in numbers of larvae in this organ at 14 dpi between the two species.

We found a similar number of *Toxocara* spp. in the MLN and lungs at 14 dpi and it is therefore proposed that the infectivity of *T. cati* in the pig host is equal to that of *T. canis*. However, this is difficult to confirm with certainty as not all organs were examined and only two time points investigated. Later in the infection course, there was a tendency for more *T. cati* in the MLN whereas most *T. canis* were found in the lungs, but differences were not significant. There is therefore a need for further studies where pigs are necropsied both at an earlier time point and at more regular time intervals during the infection period to confirm these findings. The recovery rate of larvae was lower at 31 dpi compared to 14 dpi, suggesting that larvae are redistributed within the host body with time and/or are degraded by the immune response (25).

As previous reported we found that *T. cati* can migrate to the brain of a larger animal implying that this parasite also might be involved in NT described in humans (34). Although we recovered slightly higher numbers of *T. canis* larvae from the brains than *T. cati*, the difference was not statistically significant, and further studies are needed to evaluate if *T. canis* larvae have a higher affinity for the brain tissue compared with *T. cati*. In mice, *T. canis* lead to about 10 times more differentially transcribed genes as compared to *T. cati* but both species may cause neurological symptoms and behavioural changes (13,40). No *Toxocara* spp. larvae were recovered from the eyes of the pigs confirming that OLM is a rare event in pigs infected with high infection doses (27,30,31,34).

Infections with both *Toxocara* species gave rise to white spots on the livers and kidneys at both time points in accordance to previous experimental infections studies in pigs with *T. canis* (25–27,30,31), and *T. cati* (26,34,41). The median total number of liver white spots on 14 dpi were 486 (*T. canis*) and 493 (*T. cati*) (Table 2) and similar to previous findings for *T. canis* using a similar infection dose (27,30). Unfortunately, Roneus (26) did not quantify the white spots but noted that these were more conspicuous for *T. canis*. This agrees with our findings of more lymphonodular white spots at 14 dpi giving the livers of *T. canis* infected pigs a very rough surface. This may be the reason why more liver white spots were observed later in the infection for *T. canis* compared to *T. cati* infected pigs, in accordance with (26), since lymphonodular white spots take longer time to heal, as also observed for *A. suum* (42). However, in general, a marked decrease in liver white spots is observed with time, in particular, for *T. cati* infected pigs (25–27,30,31,34).

In accordance with previous studies liver granulomas were observed in *T. canis* infected pigs (25,26,31). In contrast no granulomatous reaction was found in the livers of *T. cati* infected pigs but it cannot be excluded that granulomas may have been present in other sections. Indeed, granulomas in the liver of *T. cati* infected pigs that were similar to the ones found in *T. canis* infected pigs have been described (26). Our results support earlier studies that both infections cause lung and MLN granulomas (25,31,34,41). Two studies also observed giant cells in MLN of *T. cati* infected pigs, supporting our observations (34,41). While the finding of liver fibrosis in both infections confirms earlier studies (25,26,31,41), the presence of lung fibrosis has not previously been described in *T. cati* infected pigs. Our results indicate fibrosis may occur in MLN of *T. canis* infected pigs and as previously been described in the gastrosplenic lymph nodes (25). No previous study has examined fibrosis in lymph nodes of *T. cati* infected pigs and our results indicate that the infection does not cause MLN fibrosis.

We found that both infections caused early systemic and tissue eosinophilia in liver, lungs and MLN as previously described by other authors (25,26,31,34,41). We also observed that both infections can cause focal consolidation in the lung as observed in *T. canis* infected pigs (31).

## Conclusions

*T. canis* is commonly assumed to be the main causative agent of toxocarosis. However, we observed that *T. cati* overall had a similar migration pattern in pigs as *T. canis* and likewise induced systemic eosinophilia, white spot formation on livers and kidneys and severe histopathological changes. In addition, the study proved that *T. cati* can cause NT in a larger mammal. This study therefore emphasizes the need for further studies on the importance of *T. cati* as a zoonotic agent (15), particularly its role in NT.

## Acknowledgements

The authors would like to thank Claudia Böhm, Sonja Wolken, and Christina Strube, University of Veterinary Medicine Hannover, Germany, for providing eggs for the infection of pigs. This work was funded by the University of Copenhagen, Veterinary Parasitology Research Group. Ayako Yoshida was supported by a grant from the Japan Society for the Promotion of Science (S 2213 and 21K06995).

## Author contributions

PN, AY, PS and SMT conceived and designed the experiment. PN, AY, CSP, SMT and PS conducted the infection study and TTW and PSL analysed the histological samples. CSP conducted the statistical analysis. CSP and PN drafted the manuscript. All authors critically revised and approved the final manuscript.

## About the author

Dr. Poulsen was a master student at the Department of Veterinary and Animal Sciences, University of Copenhagen, during the period of the study. He investigated the zoonotic potential of *Toxocara* spp. and evaluated if serodiagnostics could be used to differentiate the infection type. Dr. Poulsen is currently a postdoc working on host-microbial interactions in allergic diseases.

