## Supplementary Data for "Migratory pattern of zoonotic *Toxocara cati* and *T. canis* in experimentally infected pigs"

**Appendix**

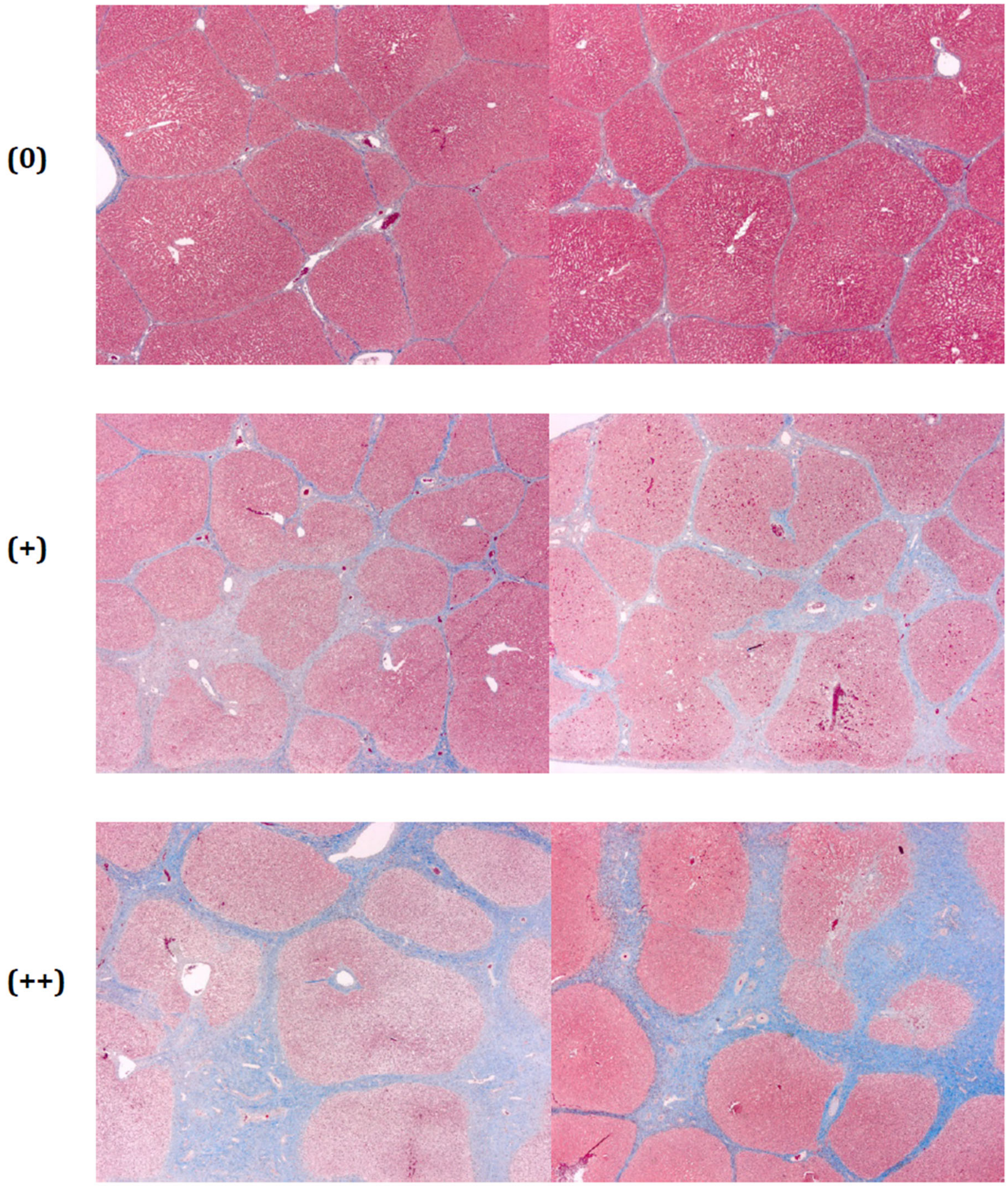


**Appendix Figure 1.** Categorization of liver fibrosis (20X): non-existent (0), moderate (+), or massive (++). Masson trichrome stains fibrous tissue light blue.

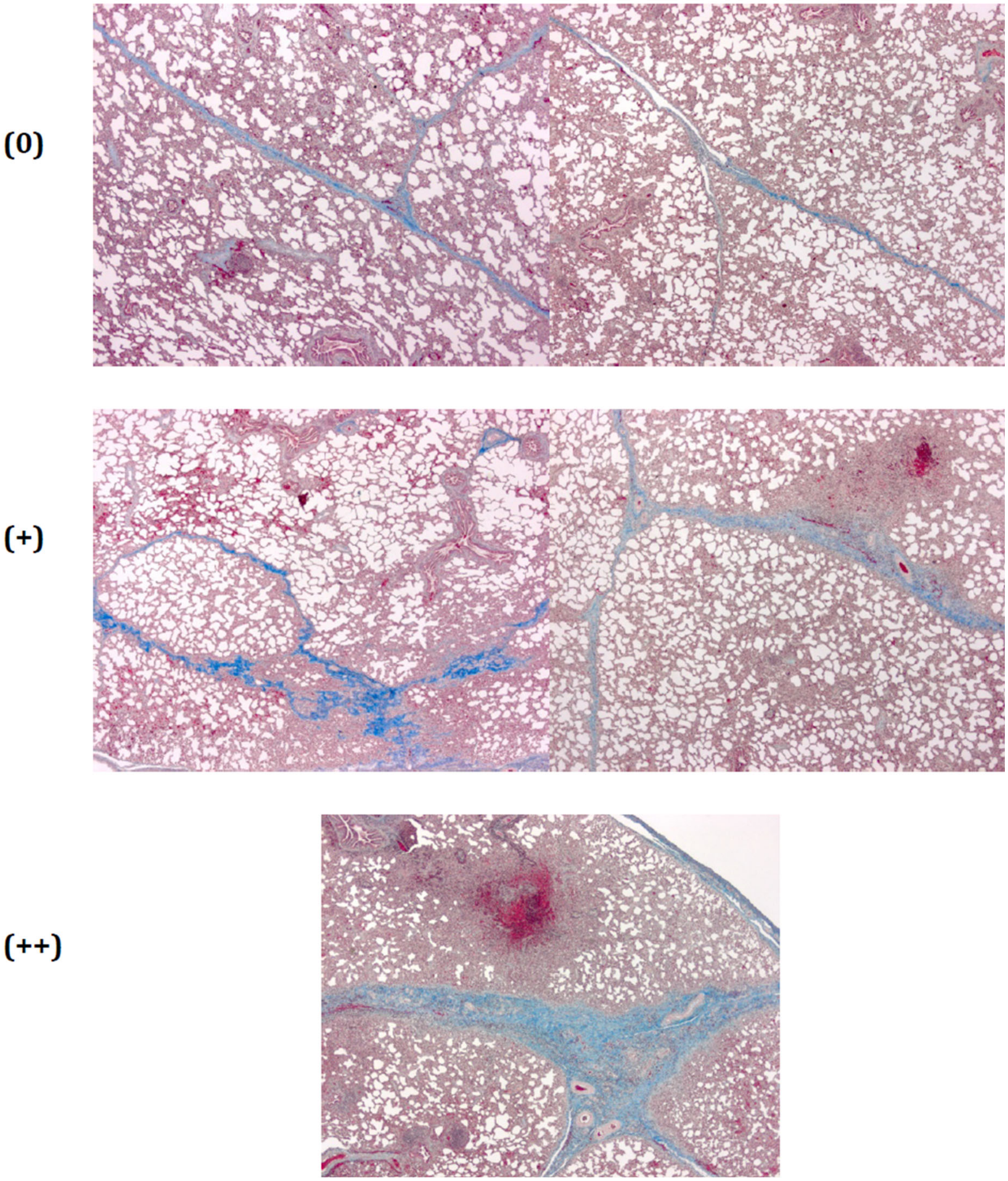


**Appendix Figure 2.** Categorization of lung fibrosis (20X): non-existent (0), mild (+), or massive (++). Masson trichrome stains fibrous tissue in the interlobular septa light blue.

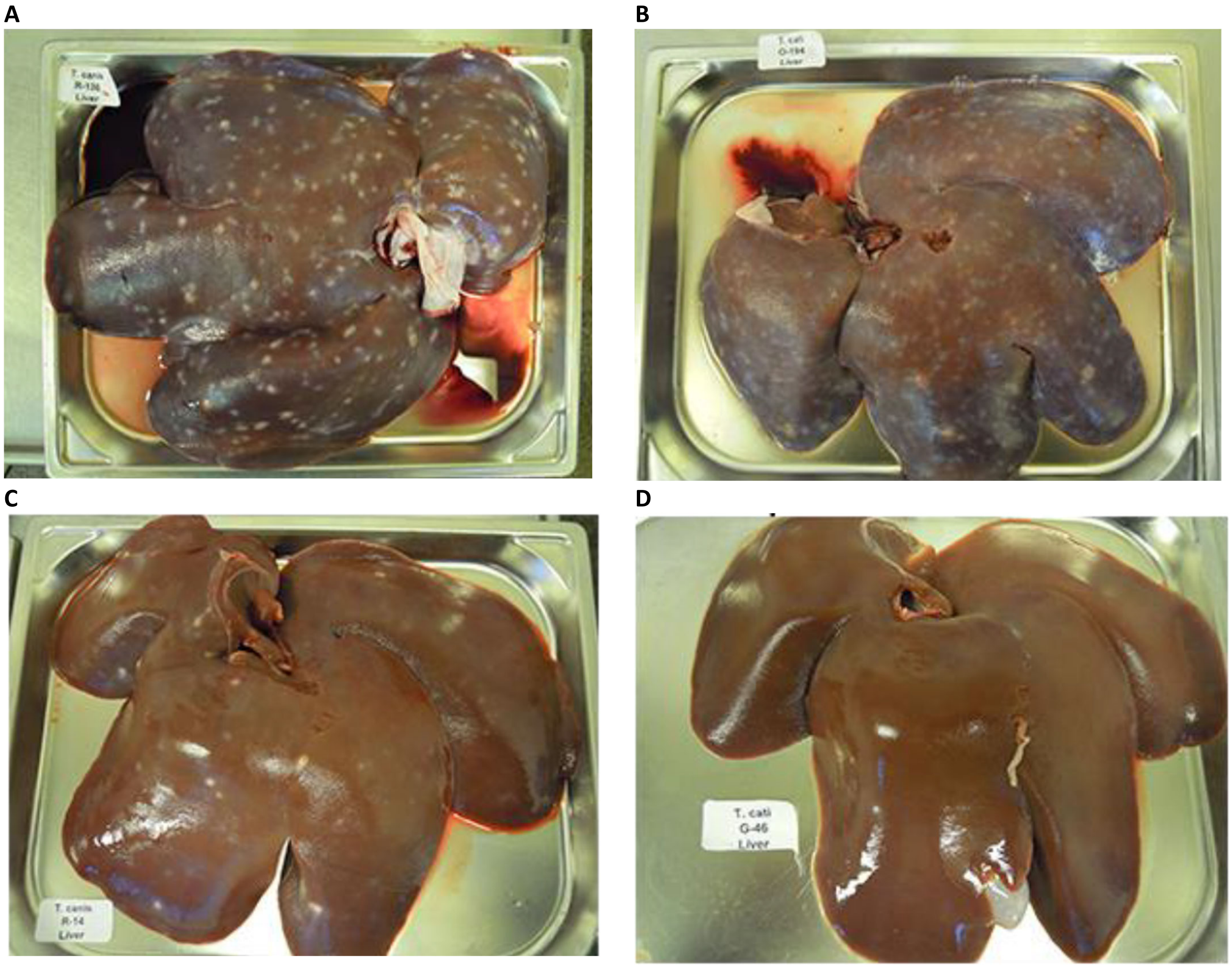


Appendix Figure 3. Representative livers of pigs infected with Toxocara spp. A and B) *T. canis* and *T. cati* infected pigs (50,000 eggs) at day 14 dpi., respectively. C and D) *T. canis* and *T. cati* infected pigs (10,000 eggs) at day 31 dpi., respectively.

Appendix Table 1. Histological evaluation of sections from indicated organs from pigs infected with 50,000 and 10,000 Toxocara spp. eggs or uninfected controls on day 14 and 31 days post infection (DPI), respectively

| Pig Id | DPI | Infection | Eosinophila | | | Fibrosis | | | Necrosis | | Granulomas | | | Emphysema |
|---|---|---|---|---|---|---|---|---|---|---|---|---|---|---|
| Scoring system | | | <25 (0), 25-50 (+), 50-100 (++), >100 (+++) | | | Nil (0), Moderate (+), Massiv (++) | Nil (0), mild (+), massive (++) | Nil (0), Present (+) | Nil (0), Present (+) | | Nil (0), 1-2 (+), 3-4 (++), 5> (+++) | | | Nil (0), Present (+) |
| Organ type | | | Liver | Lung | Lymph node | Liver | Lung | Lymph node | Liver | Lung | Liver | Lung | Lymph node | Lung |
| 42103-14-Y32 | 14 | T. canis | +++ | +++ | +++ | ++ | 0 | 0 | + | + | 0 | ++ | 0 | 0 |
| 42103-1-R136 | 14 | T. canis | +++ | +++ | +++ | ++ | 0 | 0 | + | 0 | 0 | 0 | 0 | 0 |
| 42103-3-Y33 | 14 | T. canis | +++ | +++ | +++ | ++ | 0 | 0 | 0 | 0 | 0 | + | + | 0 |
| 42103-5-R137 | 14 | T. canis | +++ | +++ | +++ | ++ | + | 0 | + | + | ++ | ++ | 0 | 0 |
| 42103-6-Y31 | 14 | T. canis | +++ | +++ | +++ | + | 0 | 0 | 0 | 0 | 0 | 0 | 0 | 0 |
| 42103-8-R138 | 14 | T. canis | ++ | +++ | +++ | 0 | + | + | 0 | + | 0 | + | + | 0 |
| 42103-10-B47 | 14 | T. cati | +++ | +++ | +++ | + | + | 0 | 0 | + | 0 | ++ | 0 | 0 |
| 42103-13-O193 | 14 | T. cati | +++ | +++ | +++ | 0 | 0 | 0 | 0 | 0 | 0 | 0 | + | 0 |
| 42103-4-O195 | 14 | T. cati | +++ | +++ | +++ | + | ++ | 0 | 0 | + | 0 | ++ | + | 0 |
| 42103-9-B48 | 14 | T. cati | + | +++ | +++ | 0 | 0 | 0 | 0 | 0 | 0 | + | + | 0 |
| 42103-2-O194 | 14 | T. cati | ++ | +++ | +++ | + | + | 0 | 0 | 0 | 0 | + | +++ | 0 |
| 42103-11-W53 | 14 | Control | 0 | 0 | ++ | 0 | 0 | 0 | 0 | 0 | 0 | 0 | 0 | 0 |
| 42103-12-W56 | 14 | Control | 0 | 0 | + | 0 | 0 | 0 | 0 | 0 | 0 | 0 | 0 | 0 |
| 42103-15-W55 | 14 | Control | 0 | 0 | +++ | 0 | 0 | 0 | 0 | 0 | 0 | 0 | 0 | 0 |
| 42103-16-W57 | 14 | Control | 0 | 0 | + | 0 | 0 | 0 | 0 | 0 | 0 | 0 | 0 | 0 |
| 42103-17-W54 | 14 | Control | 0 | 0 | + | 0 | 0 | 0 | 0 | 0 | 0 | 0 | 0 | + |
| 42103-7-W51 | 14 | Control | 0 | 0 | + | 0 | 0 | 0 | 0 | 0 | 0 | 0 | 0 | 0 |
| 41691-13-R18 | 31 | T. canis | 0 | + | +++ | 0 | + | 0 | 0 | 0 | 0 | 0 | 0 | 0 |
| 41691-15-R20 | 31 | T. canis | 0 | ++ | +++ | + | + | 0 | 0 | 0 | 0 | + | 0 | 0 |
| 41691-1-T15 | 31 | T. canis | +++ | 0 | +++ | ++ | 0 | 0 | 0 | 0 | +++ | 0 | ++ | + |
| 41691-5-R16 | 31 | T. canis | 0 | 0 | +++ | 0 | + | 0 | 0 | 0 | 0 | + | 0 | 0 |

| | | | | | | | | | | | | | | |
|---|---|---|---|---|---|---|---|---|---|---|---|---|---|---|
| 41691-6-R14 | 31 | T. canis | 0 | +++ | +++ | 0 | 0 | 0 | 0 | 0 | 0 | 0 | 0 | 0 |
| 41691-7-R19 | 31 | T. canis | 0 | + | +++ | 0 | 0 | 0 | 0 | 0 | 0 | 0 | + | 0 |
| 41691-8-R12 | 31 | T. canis | +++ | 0 | +++ | ++ | 0 | 0 | + | 0 | 0 | 0 | 0 | 0 |
| 41691-10-G47 | 31 | T. cati | 0 | 0 | +++ | 0 | 0 | 0 | 0 | 0 | 0 | 0 | + | 0 |
| 41691-11-G41 | 31 | T. cati | 0 | 0 | +++ | 0 | + | 0 | 0 | 0 | 0 | 0 | ++ | 0 |
| 41691-12-G45 | 31 | T. cati | 0 | 0 | +++ | 0 | 0 | 0 | 0 | 0 | 0 | 0 | + | 0 |
| 41691-16-G44 | 31 | T. cati | 0 | 0 | ++ | 0 | 0 | 0 | 0 | 0 | 0 | 0 | ++ | 0 |
| 41691-18-G43 | 31 | T. cati | 0 | 0 | +++ | 0 | 0 | 0 | 0 | 0 | 0 | 0 | 0 | 0 |
| 41691-2-G42 | 31 | T. cati | 0 | +++ | +++ | 0 | + | 0 | 0 | 0 | 0 | + | 0 | 0 |
| 41691-9-G46 | 31 | T. cati | 0 | 0 | +++ | 0 | 0 | 0 | 0 | 0 | 0 | 0 | 0 | 0 |
| 41691-14-R21 | 31 | Control | 0 | 0 | ++ | 0 | 0 | 0 | 0 | 0 | 0 | 0 | 0 | 0 |
| 41691-17-R22 | 31 | Control | $ | $ | 0 | $ | $ | 0 | $ | $ | 0 | $ | 0 | $ |
| 41691-3-R24 | 31 | Control | 0 | 0 | ++ | 0 | 0 | 0 | 0 | 0 | 0 | 0 | 0 | 0 |
| 41691-4-R23 | 31 | Control | 0 | 0 | ++ | 0 | 0 | 0 | 0 | 0 | 0 | 0 | 0 | 0 |

$ Section not available
